# Intraventricular infusion to circumvent the blood-brain barrier to gemcitabine

**DOI:** 10.64898/2026.05.01.722145

**Authors:** Bruno Chauffert, Antoine Galmiche, Christophe Louandre, Bernard Royer, Michel Simonet, Nelly Guilain, Fabien Rech, Pierre Simonet, Maxime Sibert, Ahmed Abdaoui, Alexandre Cau, Mathieu Boone, Jacques Beaurain

## Abstract

The poor prognosis of brain tumors, including IDH-wild-type glioblastoma (GB), as well as brain and leptomeningeal metastases, is partly related to the blood–brain barrier (BBB), which limits the delivery of hydrophilic anticancer drugs to the tumor site and surrounding brain parenchyma. Early studies using vital dyes demonstrated that intracranial injection could bypass the BBB in cats. We confirmed that, in guinea pigs, the vital dye Bleu Patente V diffused efficiently into the brain after a bolus intracranial injection, whereas the brain remained unstained after intravenous administration. Similarly, brain concentrations of the hydrophilic anticancer drug gemcitabine were significantly higher following intracranial injection than after intravenous administration. Consistent with these findings, Bleu Patente penetrated deeply into the cerebral cortex of sheep after a 24-hour intraventricular infusion. At the end of a 24-hour intraventricular infusion of 20 mg gemcitabine in sheep, mean gemcitabine concentrations reached 1,415 µg/L in cerebrospinal fluid and 850 µg/kg in brain tissue. These concentrations exceeded the IC90 values of gemcitabine for A172, U87-MG, and U118-MG human glioblastoma cell lines, as determined in vitro after 24 hours of incubation. We hypothesize that Bleu Patente dye and gemcitabine circumvent the blood-brain barrier (BBB) by utilizing the glymphatic system. Tolerance of a single 24-hour intraventricular infusion of gemcitabine at doses of 5, 10, and 20 mg was good. Taken together, these encouraging preclinical results support the resumption of Phase I clinical trials evaluating intraventricular infusion of gemcitabine in patients with refractory primary or secondary brain tumors.

## INTRODUCTION

The poor prognosis of brain tumors, including IDH-wild-type glioblastoma (GB), as well as brain and leptomeningeal metastases, is partly attributable to the blood–brain barrier (BBB), which restricts the delivery of hydrophilic anticancer drugs to the tumor site and to the surrounding brain parenchyma infiltrated by malignant cells. Consequently, circumventing the BBB remains a major therapeutic challenge. Several active clinical strategies are currently being evaluated to bypass the BBB, including focused ultrasound, nanovesicle-based delivery systems, and membrane transporters (1, 2). Early experimental studies demonstrated that intraventricular administration of vital dyes could bypass the BBB in animal models. Trypan Blue (3) and Bromophenol Blue (4) failed to stain brain parenchyma when administered intravenously in cats, whereas deep parenchymal penetration was observed following intraventricular injection. We reproduced and extended these historical experiments in guinea pigs and sheep using Bleu Patente V dye (BPD) as a hydrophilic vital dye and gemcitabine as a hydrophilic anticancer drug.

Gemcitabine (GEM) is a widely used cytotoxic agent with proven clinical efficacy in a broad range of solid tumors, including lung, breast, and pancreatic cancers, but not in gliomas. Nevertheless, several studies have demonstrated significant in vitro activity of GEM against human GB cell lines, whereas clinical translation using the intravenous route has so far been unsuccessful (5). Similarly, GEM is less effective against cerebral and leptomeningeal metastases than against extracerebral tumor sites. This discrepancy between strong *in vitro* efficacy and limited clinical activity in brain tumors is attributable to the restrictive BBB permeability to GEM (6).

In this context, we developed a protocol for a 24-hour intraventricular infusion of gemcitabine designed to circumvent the BBB in sheep and enhance its antitumor activity against primary and metastatic brain tumor cells. We evaluated the tolerance of intraventricular GEM administered at escalating doses in sheep, establishing the feasibility of this delivery strategy.

## MATERIAL AND METHODS

### Cell culture and reagents

Human GB cell lines A172, U87-MG and U118-MG were purchased from American Type Culture Collection (ATCC) (LGC Standards, Strasbourg, France) and cultured in Dulbecco Modified Eagle Medium (DMEM) supplemented with 10% fetal calf serum (Jacques Boy, Reims, France), 2 mM glutamine and penicillin / streptomycin. Gemcitabine and cytosine arabinoside were purchased from Sigma Aldrich for *in vitro* assays. Gemcitabine for clinical use was used for animal studies (Gemcitabine Arrow, 40 mg/ml). Bleu Patente V dye (BPD, CI 42051, sulfan blue, MW 566) for clinical use was obtained from Guerbet (Villepinte, France).

### In vitro cytotoxicity of GEM

Cells were seeded and cultured for 48 h before an exposure for 1 hour or 24 h to GEM. After rinsing wells, cells were cultured again without any drug for 72 h. Surviving cells that adhered to the well bottom were fixed with ethyl alcohol then stained with Crystal Violet (1 %) for 10 min. After three steps of water rinsing, Crystal Violet was eluted with 1% SDS solution. Optical density (OD) was read at 595 nm. Cell survival (CS) was calculated using the formula: CS = mean OD in treated wells/mean OD in control wells X 100. IC50 and IC90 were the concentration that inhibit 50 or 90 % of cell survival on graphs, respectively.

### Immunoblot assays

Total protein extracts were obtained after cell lysis in RIPA buffer. Standard immunoblot protocols were used for their analysis by immunoblot. The antibodies that we used are: anti-phospho-CHK1 Ser345 (Cell signaling, 2348), anti-CHK1 (Abcam, ab32531), anti-γ-H2AX (7631, Cell signalling), anti-phospho-AKT Ser473 (Cell signaling, 4060), anti-AKT (9272, Cell signalling), anti-ERK1/2 (Cell signaling, 9102), anti-phospho-ERK1/2 Thr202/Tyr204 (Cell signaling, 9101), anti-Lamin B1 (Santa Cruz, sc-374015), anti-β-actin (Sigma-Aldrich, A5441).

### ELISA assays

ELISA experiments were conducted on cell culture supernatants. The human uPAR ELISA kit was purchased from R&D Bio-techne (reference DUP00), and experiments were performed according to the manufacturer’s protocol.

### Cytokine array

The Sheep Cytokine Array was purchased from AssayGenie (reference SARB0092) was used in order to monitor the protein levels of 18 cytokines in the CSF: CCL5, DCN, FRZB, IFNG, IL-4, IL13, IL17A, IL1A, IL1B, IL21, IL4, LIF, SPINK5, TNF, TNFA, TNFSF2, VEGF, VEGFA.

### In silico analysis of cell lines sensitivity to GEM

Information regarding the sensitivity of glioma and other cancer cell lines to GEM was retrieved from The Cancer Cell Line Encyclopedia (7, 8), using the web portal: https://www.cancerrxgene.org/compound/Gemcitabine/135/overview/ic50 (accessed on the 13th of January 2025).

### Intravenous, intracerebral, and intraventricular drug administration in animals

To investigate the organ distribution of hydrophilic molecules following intravenous (IV) administration in a small animal model, adult male guinea pigs (400–500 g) received a slow IV injection via a surgically exposed jugular vein. A total volume of 1 mL, consisting of Patent Blue dye (0.5 mL) and gemcitabine (20 mg in 0.5 mL), was administered over 1 minute. Animals were anesthetized with an intraperitoneal injection of ketamine (100 mg/mL; Virbac, France; 1 mL per animal). For intracerebral administration, the scalp was incised and the skull was perforated by careful manual rotation of a 21-gauge needle. The drilling site was located 2 mm to the left of the midline along the line connecting the posterior margins of the orbits. Using a 24-gauge needle, a total volume of 0.2 mL (0.1 mL Patent Blue dye and 0.1 mL gemcitabine, corresponding to 4 mg) was injected perpendicularly to the skull at a depth of 3 mm over 1 minute. Animals were maintained under anesthesia for 30 minutes, with supplemental IP ketamine administered as required, before brain removal under deep anesthesia to minimize the risk of post-mortem disruption of the blood–brain barrier. A complete autopsy was performed, including photographic documentation, and organ samples were frozen at −20 °C prior to gemcitabine quantification.

Because prolonged intraventricular infusion is not feasible in guinea pigs, additional experiments were conducted in sheep, a large-animal model with a body weight comparable to humans. Female sheep aged 3–4 years (45–60 kg) underwent catheter placement into the left lateral ventricle. Anesthesia was induced by intravenous administration of a combination of tiletamine, zolazepam, medetomidine, and lidocaine, and maintained with 2% isoflurane. Throughout the procedure, animals received intravenous Ringer’s lactate supplemented with lidocaine for analgesia. A trephine hole (2 mm diameter) was manually drilled in the skull, located 10 mm posterior to the line connecting the posterior margins of the orbits and 5 mm lateral to the midline. A single-end-hole ventricular catheter (1.5 mm) mounted on a metal mandrel was inserted vertically through the trephine hole to a depth of 25 mm from the skull surface. Correct intraventricular positioning was confirmed by aspiration of cerebrospinal fluid (CSF). Catheter was secured to skull by cyanoacrilyc glue and tunneled under the skin of the skull to the occipital region.

Bleu Patente V (2.5%, 2 mL) was diluted in 50 mL of sterile saline and infused into the awake sheep over 24 hours using an elastomeric pump delivering a constant flow rate of 2 mL/h (D-Fuzz pump, Androme Medical, France). Animals were subsequently re-anesthetized for brain removal. CSF was collected by suboccipital puncture at a site distant from the ventricular catheter. A complete autopsy with photographic documentation was performed. The same protocol was used for intraventricular infusion of gemcitabine (20 mg in 50 mL saline over 24 hours). Organ samples were collected and stored at −20 °C until gemcitabine assay.

All animal experiments were conducted by a senior veterinary doctor. Animals were anesthetized for each experiment and euthanized by lethal injection of barbiturates at the time of sampling. In accordance with the 2010/63/UE European directive and French regulations at the date of the animal experimentation, every effort was made to minimize animal suffering, and the study adhered to widely accepted principles for animal care and use.

### GEM quantification in biological samples

The frozen tissues were weighted, then a mixture of tetrahydrouridine (THU - a cytidine deaminase inhibitor that prevents a potential GEM degradation in vitro – Alsachim, Illkirch-Graffenstaden, France) (25 µg/mL) and acetonitrile (50/50 v/v) was added (600 µl for about 100 mg of tissue). Tissues were then submitted to an ultrasonic homogenizer (Pulse 150, Benchmark Scientific, USA) then centrifugated at 20 000 g. Samples were diluted to 1/10^th^ and then a mixture of internal standard (2N^15^, C^13^ GEM, Alsachim, Illkirch-Graffenstaden, France) with acetonitrile (50/50 v/v) was added to 200 µL of supernatant, vortexed, and evaporated to dryness under nitrogen. The residues were reconstituted with water, then analysed using LC/HRMS. In brief, an Atlantis T3 column (Waters, Saint-Quentin-en-Yvelines, France) was used for chromatographic separation, and an Exploris 120 (ThermoFisher Scientific, San Jose, USA) was used for detection. Cerebral spinal fluid and urine underwent the same extraction process, except that THU was not used. The dilution factor was 1/100^th^.

### Histological examination of the brain

Whole brains of sheep were fixed with 4% paraformaldehyde for 14 days. 5 µm sections were stained with Hematein-Eosin-Saffran and examined on a Zeiss microscope.

### Statistics

IC50 and IC90 were determined using GraphPad Prism5. Student’s T test was calculated with Excel. A threshold of p = 0.05 was retained for significance.

## RESULTS

### Circumventing BBB to Bleu Patente dye by intracranial injection in animals

We compared the tissue distribution of the hydrophilic molecule Bleu Patente dye (BPD) following intravenous or intracranial administration in male guinea pigs. Animals were euthanized and autopsied 30 minutes after injection, corresponding to the time of maximal peripheral coloration (tongue and anterior paws) in intravenously treated animals. After intravenous injection, all examined organs exhibited blue coloration, with the notable exception of the brain and testicles (Figures 1A and 1B). The presence of blue dye in the gallbladder, urinary bladder, and intestinal contents indicated that BPD is eliminated through both hepatic and renal pathways. In contrast, following intracranial injection, the brain displayed intense and homogeneous blue coloration 30 minutes after administration, while peripheral organs initially remained unstained. Notably, the gallbladder, urinary bladder, and intestinal contents were also blue, indicating a rapid systemic redistribution of BPD from the central nervous system into the circulation for subsequent elimination.

**Figure 1.**
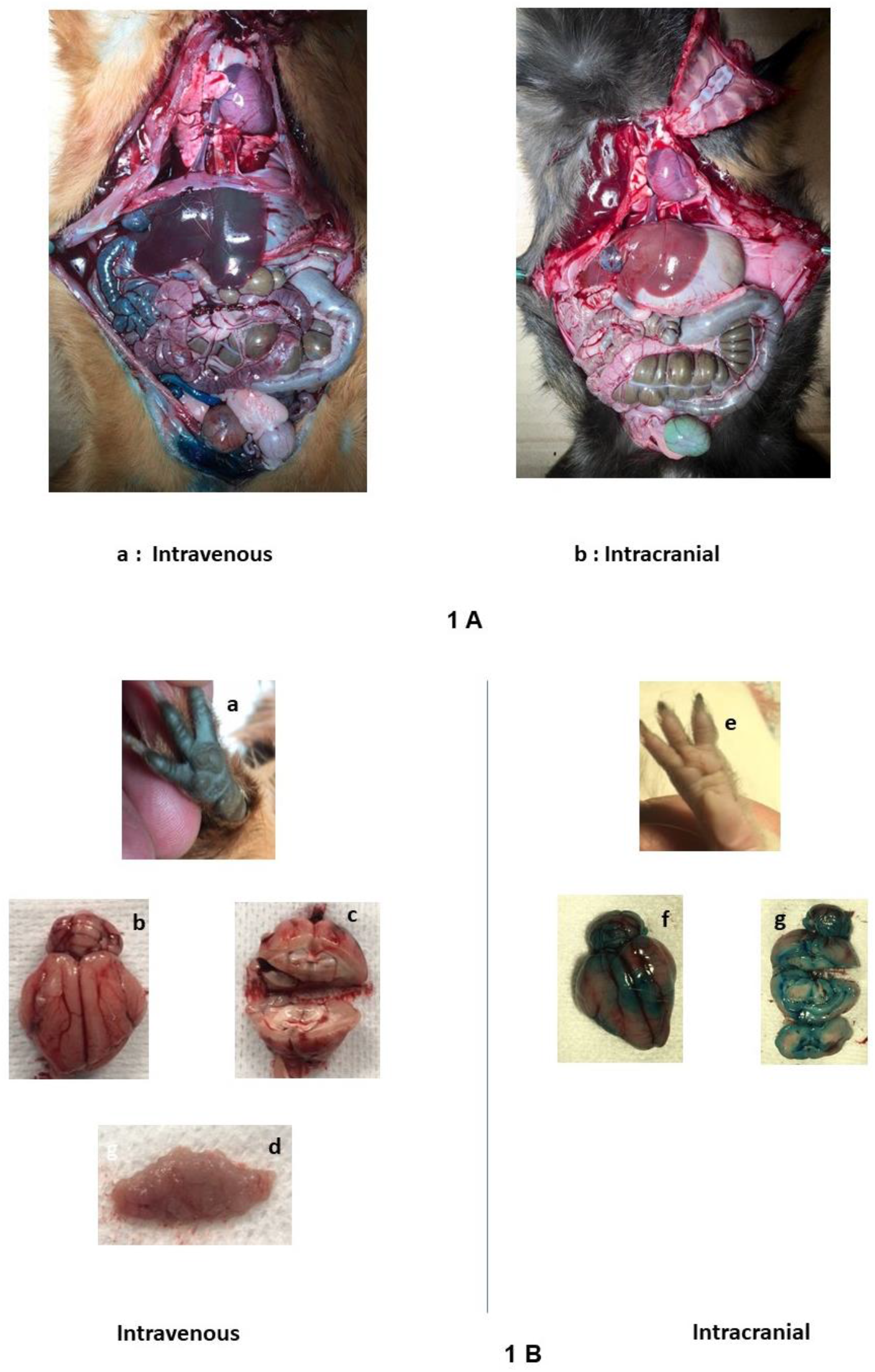
Bleu Patente dye distribution in organs after intravenous or intracranial injection in guinea pigs. Three animals received Bleu Patente dye (BPD) either by intravenous route (0.5 ml) or by an intracranial injection (0.1 ml) (Figure 1 A). Autopsy was done at 30 min when anterior palms were stained at maximum after the IV injection (Figure 1 B a). After IV injection, all organs were blue (Figure 1 A a) except brain and testicles (Figure 1 Bb, 1 Bc, 1 Bd). Content of gallbladder, intestine and bladder was blue. After intracranial injection, brain (surface: figure 1 Bf and depth: figure 1 Bg) was the only intensively stained organ. All other organs were unstained as anterior palms (Figure 1 Be, Figure 1 Ab). Content of gallbladder, intestine and bladder was blue after intracranial injection, reflecting the elimination of the dye via urine, liver and then intestine.

The diffusion of Bleu Patente dye (BPD) into brain parenchyma was subsequently evaluated in sheep after intraventricular administration. One hour after a bolus intraventricular injection, limited penetration of dye into the brain parenchyma was observed, predominantly within the periventricular regions and the adjacent peripheral gray matter (Figure 2A). In contrast, following a 24-hour continuous intraventricular infusion, BPD showed markedly deeper penetration into the cerebral gray matter, whereas staining of the hemispheric and thalamic white matter remained limited (Figure 2B). Notably, dye impregnation was more pronounced in the inferior cortical regions of the cerebral hemispheres, likely reflecting gravitational effects in sheep, which are rarely positioned horizontally. Consistent with this observation, the cortical layer of the cerebellum exhibited intense blue staining, while the underlying medulla remained unstained (data not shown).

**Figure 2.**
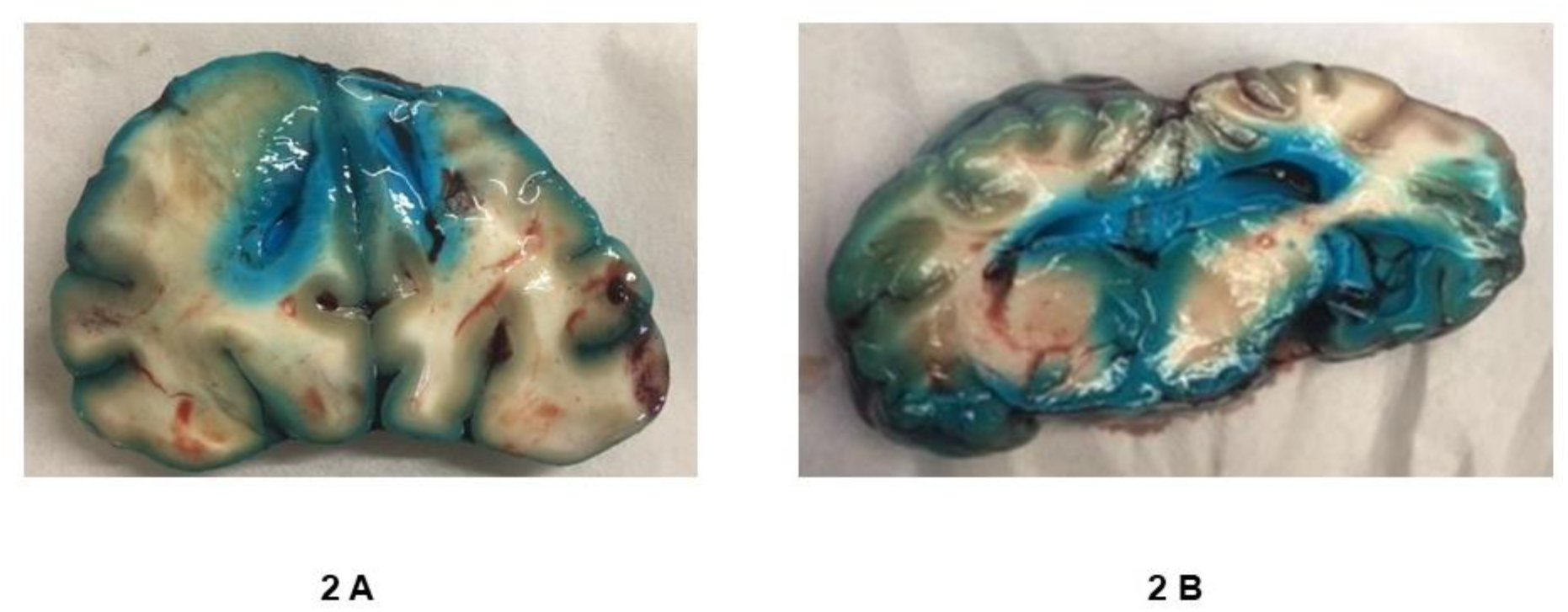
Bleu Patente distribution in brain after intraventricular infusion in sheep. Animals received 2 ml BPD either as a bolus injection into the left ventricle 1 h before autopsy (Figure 2 A) or after a 24 h infusion (Figure 2 B). Representative photographs from 2 animals by condition.

### Circumventing BBB to gemcitabine by intrathecal injection in animals

We measured GEM distribution in organs of male guinea pigs either 30 min after the IV injection of 20 mg GEM or the intracranial (IC) injection of 4 mg of GEM (Figure 3). The BBB to GEM after an IV injection was confirmed. GEM concentration (16 µg/kg) was 5.1 fold less in the brain than in the liver, kidney and lung (mean concentration 82 µg/kg, p = 0.001, Student’s t test). The barrier between blood and testicle was also confirmed. After an IV injection, GEM concentration of was 2.5 fold less in the testicle parenchyma (32 µg/kg) than in the liver, kidney and lung (82 µg/kg, p = 0.003, Student’s t test). A strong advantage of the intracranial injection over the IV route was evidenced. GEM concentration in brain was 662 µg/kg after an intracranial injection compared to 16 µg/kg after an IV injection (fold X 41, p < 0.0001 Student’s t test) despite the lower dose (4 mg by intracranial vs 20 mg by IV route). GEM was detected in the liver, kidney, lung and urine even after an intracranial injection indicating a recirculation of the drug and its elimination by liver and urine.

**Figure 3.**
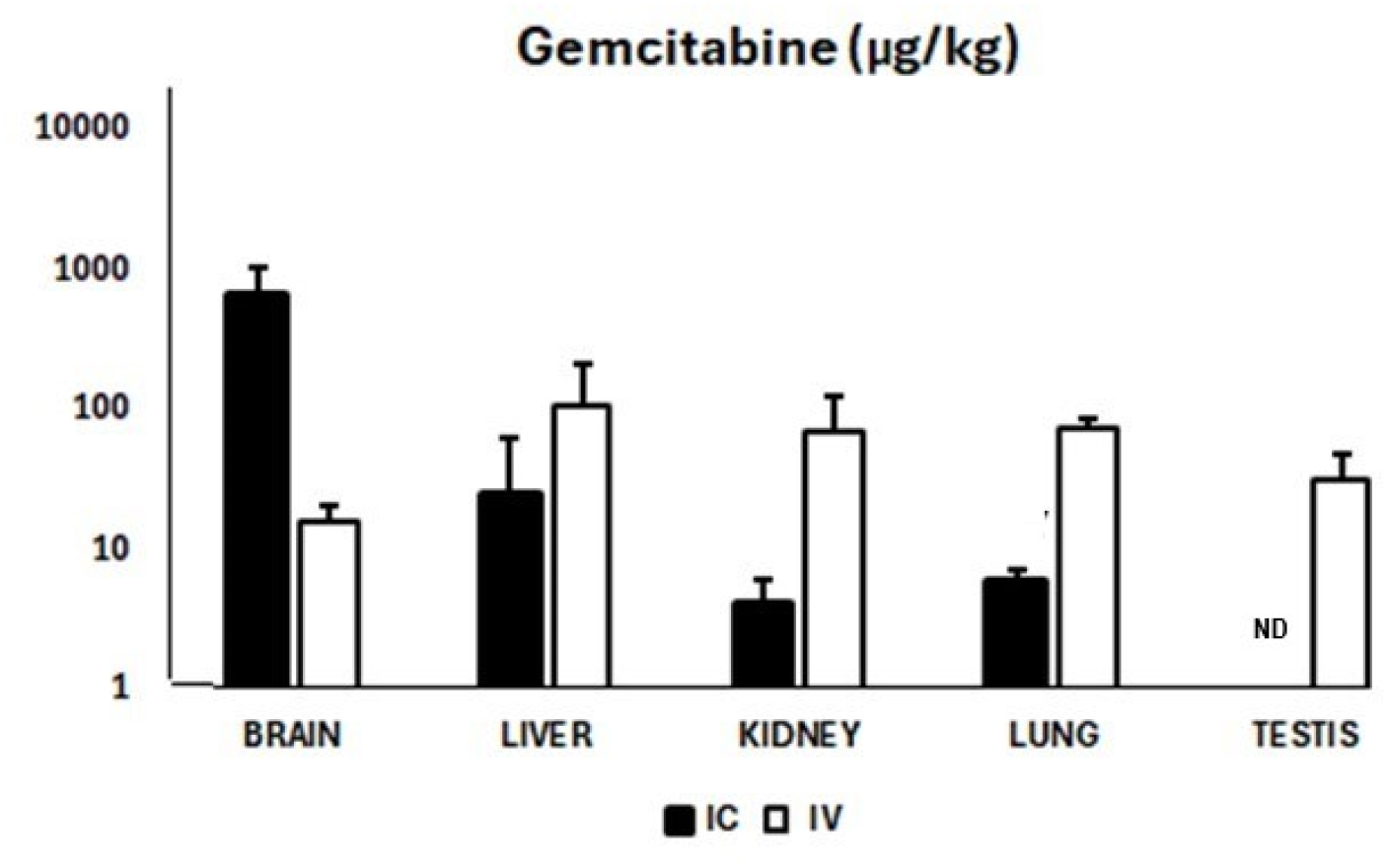
GEM distribution after an intravenous or intracranial injection in guinea pigs. GEM was assayed in the organs either after an intravenous injection of 20 mg GEM (clear bars) or an intracranial injection of 4 mg GEM (dark bars). Mean of 3 animals for each condition. ND: *not determined*.

We measured GEM distribution in cerebrospinal fluid, brain and other organs of 3 sheep at the end of a 24 h infusion of 20 mg GEM by the intraventricular route (Figure 4). Mean GEM concentration was 1 415 µg/l in CSF and 850 µg/kg in the brain. GEM was eliminated in urine whereas GEM concentration was low in liver, kidney and lung.

**Figure 4.**
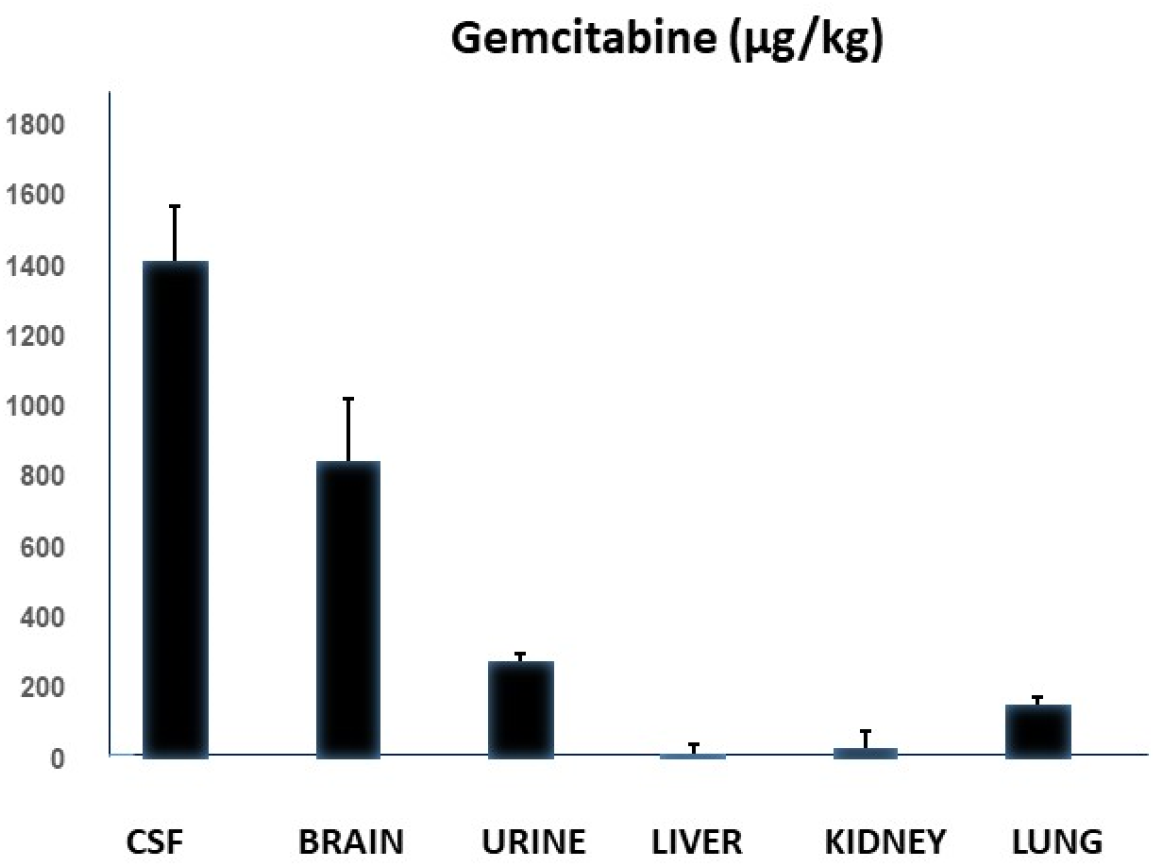
GEM distribution after an intraventricular infusion in sheep. GEM was assayed in the organs of 3 animals at the end of a 24 h intraventricular infusion of 20 mg. Mean from 8 determinations (4 by hemisphere) for the brain and 1 determination by animal for CSF, urine, liver, kidney and lung.

### Prolonged exposure increases GEM cytotoxicity against human glioblastoma cells

To evaluate whether gemcitabine (GEM) concentrations achieved in cerebrospinal fluid (CSF) and brain tissue of sheep after a 24-hour intraventricular infusion would be sufficient to kill glioblastoma (GB) cells, we performed *in vitro* cytotoxicity assays using three human GB cell lines: A172, U87-MG, and U118-MG. Cells were exposed to GEM for 1 hour or 24 hours to mimic the pharmacokinetic conditions of a bolus intravenous injection or a 24-hour intraventricular infusion, respectively. A 24-hour exposure to GEM induced a marked reduction in cell viability, with average IC50 values of 5.3 nM (1.39 µg/L), 8.7 nM (2.29 µg/L), and 7.9 nM (2.08 µg/L) in U87, U118, and A172 cells, respectively. The IC90 values for all three lines were below 100 nM (26 µg/L) (Figure 5A). In contrast, brief 1-hour exposure resulted in substantially reduced cytotoxicity: IC50 values were 139 nM (36.6 µg/L), 294 nM (77.4 µg/L), and 387 nM (101.9 µg/L) in U87, U118, and A172 cells, respectively. Under 1-hour exposure conditions, even the highest tested GEM concentration (10 µM, 2 500 µg/L) failed to eradicate the cells. Notably, GEM was markedly more active than cytosine arabinoside under comparable 24-hour treatment conditions (Figure 5A). Microscopic examination of residual adherent cells five days after 24-hour exposure to GEM at IC90 revealed cells with enlarged morphologies, appearing either flattened or with branched processes (Figure 5B), possibly suggestive of cellular senescence. To explore the mechanisms underlying GEM-induced cytotoxicity, we assessed its effects on key oncogenic signaling pathways and the DNA-damage response (DDR) via checkpoint kinase 1 (CHK1). Levels of phosphorylated ERK1/2, Protein kinase B (PKB/Akt), and CHK1 (Ser345) were analyzed by immunoblotting after 48 hours of GEM exposure (Figure 6A). GEM did not alter ERK or PKB phosphorylation but induced a concentration-dependent increase in CHK1 Ser345 phosphorylation, consistent with activation of DDR and genotoxic stress. Given the observed senescent-like morphology, we evaluated Soluble urokinase-type plasminogen activator receptor (uPAR), a recently reported senescence marker (9), in culture supernatants from GB cells transiently exposed to GEM (10 nM or 1 µM) for 24 hours and then maintained for five days in fresh medium. All three cell lines showed increased soluble uPAR levels (Figure 6B). We also assessed the nuclear lamina component lamin B1, which is downregulated in senescent cells (10). Immunoblot analysis revealed clear reductions in lamin B1 levels in U87, U118, and A172 cells (35%, 51%, and 35% reduction, respectively, after 1 µM GEM exposure; Figure 6B). However, p16 and p21 proteins were undetectable under these conditions (data not shown). So not all the biochemical markers of a definitive senescence are present in the residual cells (< 10 %).

**Figure 5.**
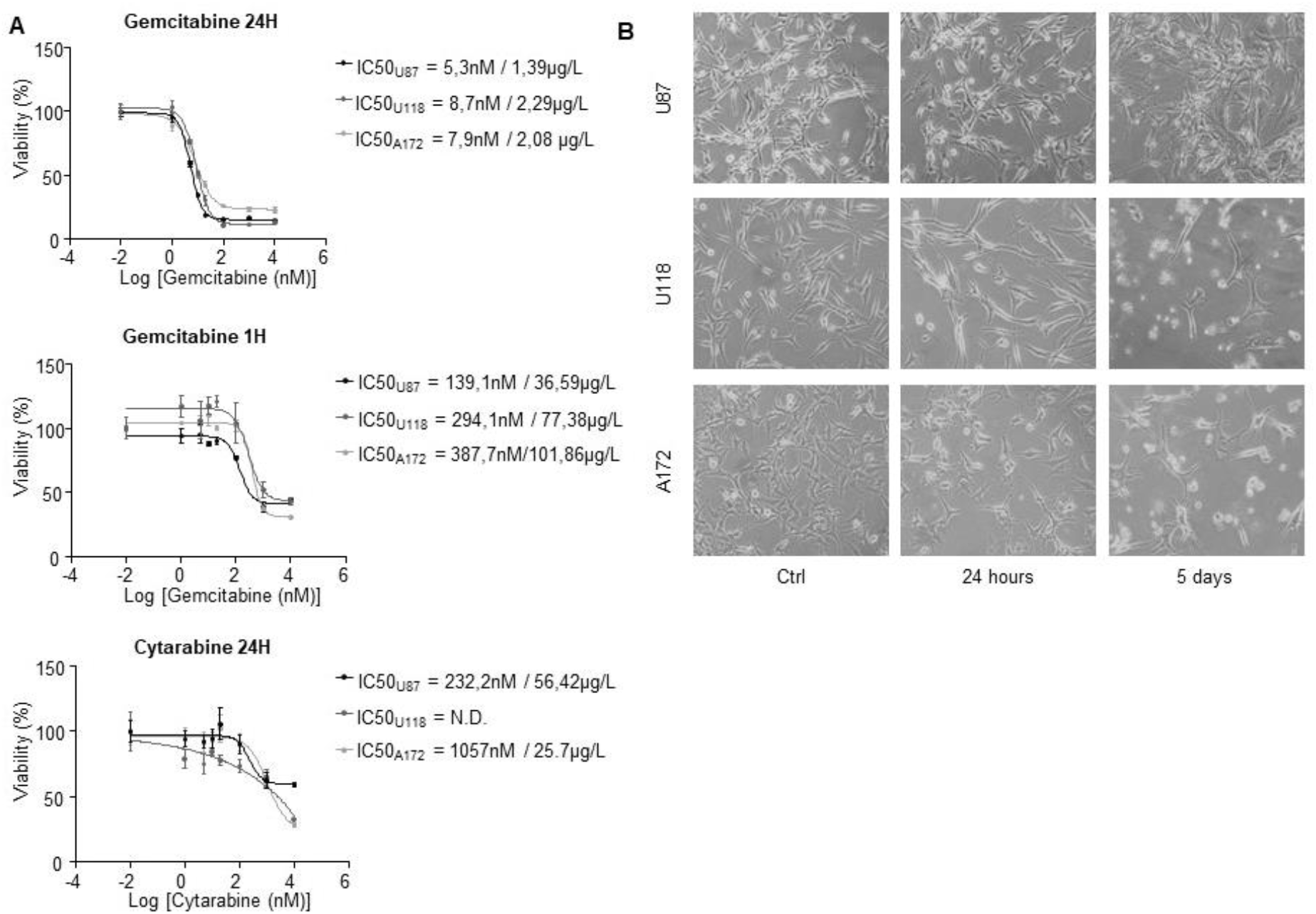
Effects of GEM exposure on the viability of GB cells. GB cells were exposed to GEM for 1h or 24 h at the indicated concentrations. After GEM exposure, cells were maintained in drug-free fresh culture medium for 72 h before a viability assay with Crystal Violet. IC50 of GEM on the A172, U87-MG and U118-MG human GB cell lines were indicated in the legend of Figure 5 A. Phase contrast microscopy images of GB cells treated for 24 h at IC 90. Cells were maintained in culture in drug free medium and observed 24 h or 5 days after GEM treatment. Note the presence of enlarged / flattened cells.

**Figure 6.**
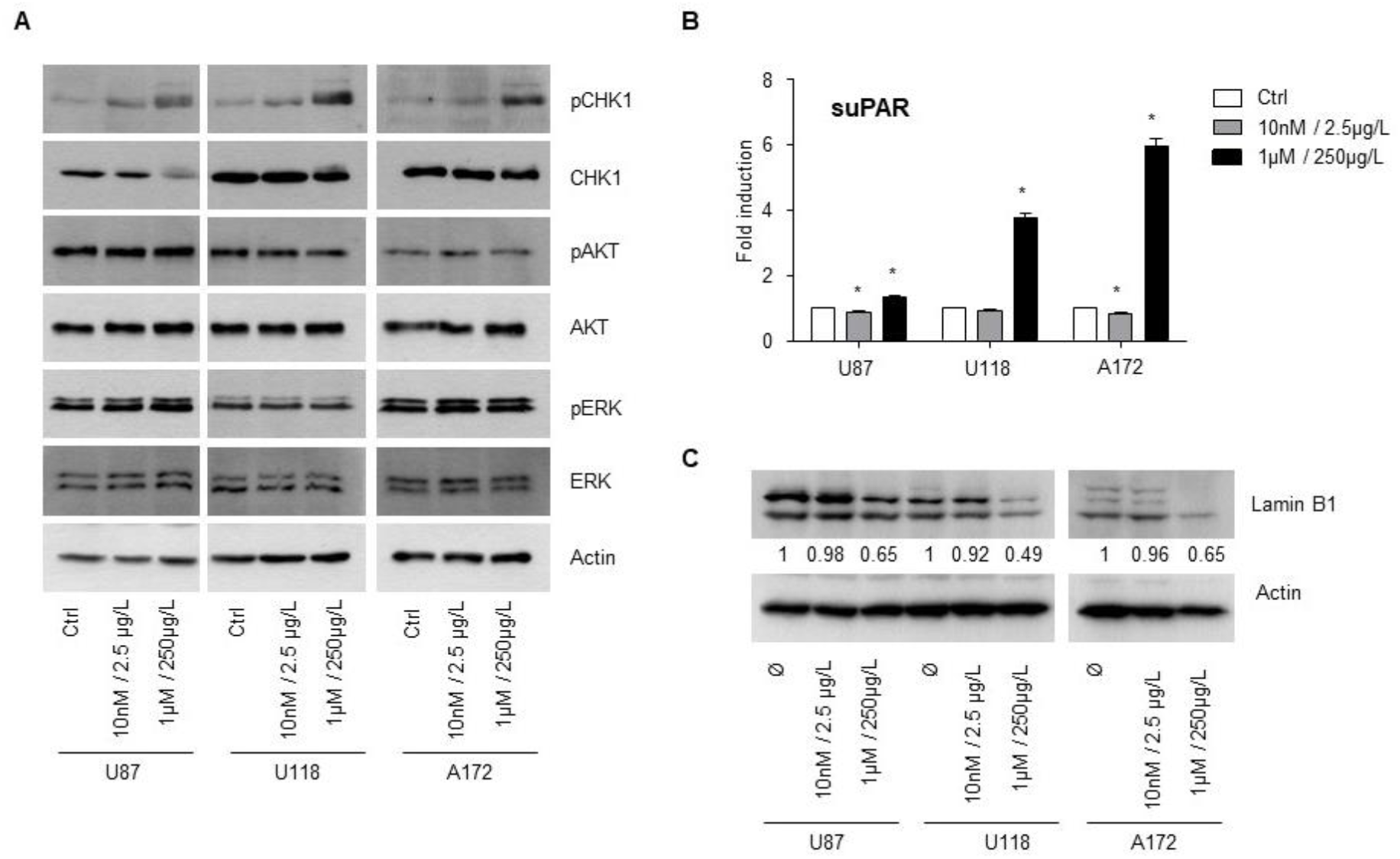
Genotoxic effects and markers of cell senescence in cells exposed to GEM. GB cells were exposed to GEM for 24h at the indicated concentrations. Culture supernatants were collected five days after a 24h of exposure to GEM at indicated concentration (Figure 6). Immunoblot analysis of the expression of the markers pCHK1/CHK1, pAKT/AKT and pERK/ERK. Actin was used as a loading control (Figure 6 A). suPAR concentrations in culture supernatants (Figure 6 B). Results are expressed as fold-induction compared to control (* indicates p <0.05 with Student’s t test comparing each experimental condition to control). Immunoblot analysis of Lamin B1 expression in GBM cells exposed to GEM (Figure 6 C).

### Assessment of neurotoxicity following 24-hour intraventricular infusion of GEM in sheep

To evaluate the potential neurotoxicity of gemcitabine (GEM) administered via 24-hour intraventricular infusion, sheep were treated with escalating doses of GEM. No clinical signs of neurotoxicity were observed at doses of 2.5 mg (n = 1), 10 mg (n = 1), or 20 mg (n = 3). Among the three animals receiving 20 mg GEM, two were euthanized for 15 days post-treatment for histological analysis, while one remains alive 12 months after treatment. Standard histological examination of the brains from the two euthanized sheep revealed no significant structural alterations (Figure 7). Analysis of cerebrospinal fluid (CSF) collected 15 days post-infusion demonstrated no major changes in cytokine profiles compared with a control animal (Figure 8). Overall, the cytokine assay did not reveal evidence of ongoing neuroinflammation. A single exception was a moderate increase in soluble Frizzled-related protein 3 (sFRP3), an inhibitor of the protumoral Wnt/β-catenin pathway, suggesting a potential modulatory effect rather than toxicity. These findings indicate that 24-hour intraventricular infusion of GEM at doses up to 20 mg is well tolerated and does not induce overt neurotoxicity in the sheep model.

**Figure 7.**
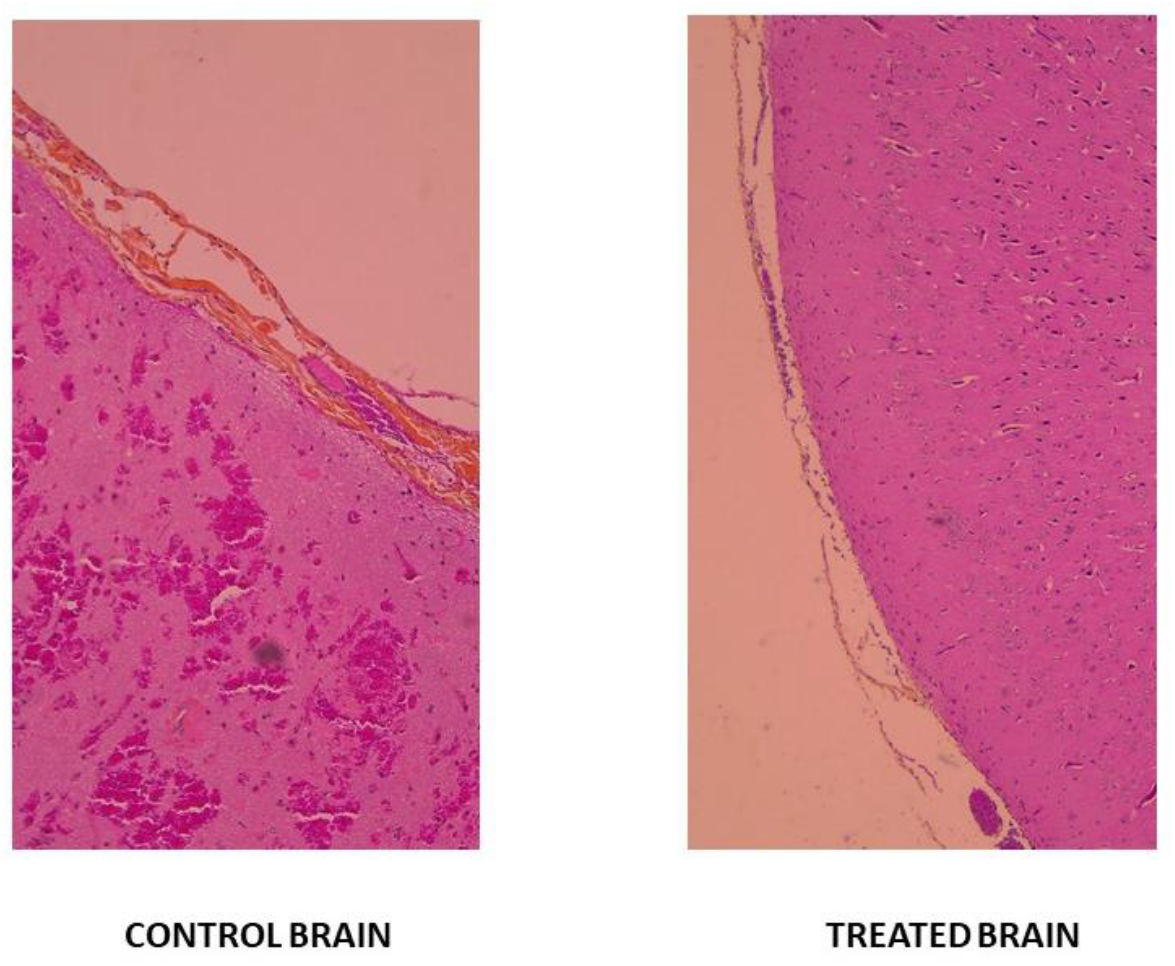
Histological examination after intraventricular GEM infusion. Two sheep received a 24 h infusion of 20 mg GEM 15 days before autopsy and brain sampling. Representative photographs did not show obvious alteration of the leptomeninges and brain in comparison with one control untreated animal.

**Figure 8.**
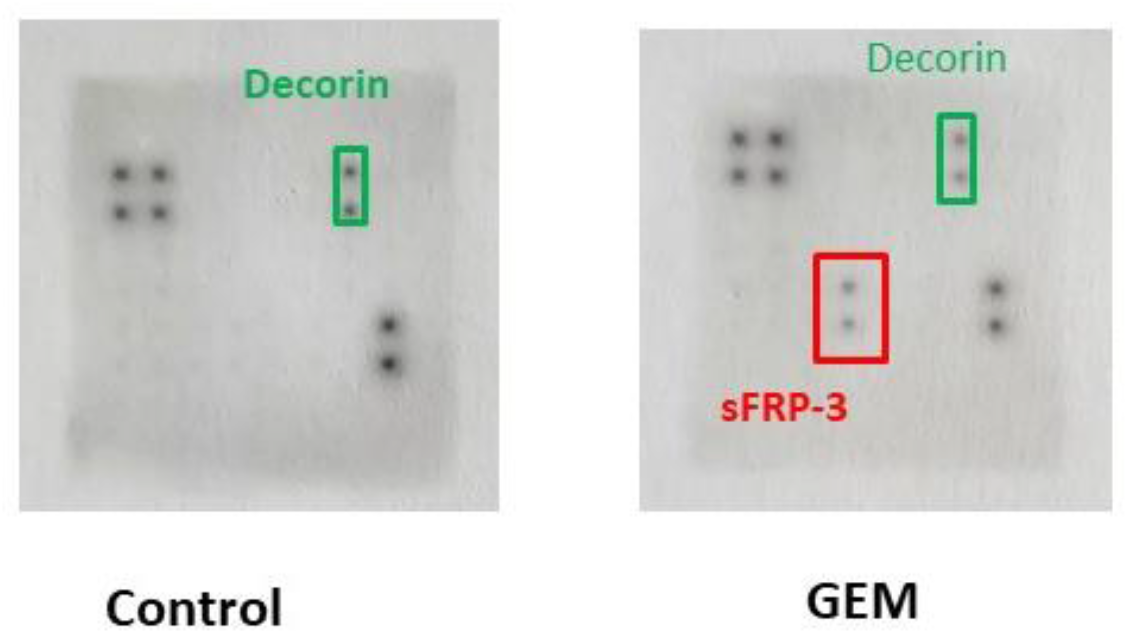
Sheep Cytokine Array. Two sheep received 20 mg of gemcitabine via a 24-hour intraventricular infusion. CSF samples were collected by occipital puncture 15 days later. No significant changes in neuroinflammatory cytokine levels were observed in treated animals compared with a control sheep.

## DISCUSSION

We confirmed the pioneering experiments of Goldman and Feldberg, which first provided evidence for the existence of the blood–brain barrier (BBB) (3,4). Following intravenous injections in cats, hydrophilic dyes did not penetrate the brain, whereas intracranial injection resulted in clear parenchymal staining. We obtained comparable results in guinea pigs after intravenous or intracranial administration of Patent Blue dye. Because prolonged intraventricular infusion is technically easier to perform in larger animals, we subsequently developed a sheep model. In this model, Patent Blue dye was detected in the cerebral cortex as early as 1 hour after a ventricular bolus injection. Dye penetration was markedly greater after a 24-hour intraventricular infusion; however, white matter remained less intensely stained than gray matter, which may be related to its higher lipid content. Similarly, we demonstrated in guinea pigs that the hydrophilic anticancer drug gemcitabine accumulates to a greater extent in the brain following intracranial administration than after intravenous injection. These findings support the conclusion that the BBB restricts the passage of hydrophilic molecules and that this barrier can be at least partially circumvented by intracranial or intraventricular delivery. Cerebrospinal fluid (CSF) and associated molecules are thought to be reabsorbed through perivascular spaces of the pial arteries, which extend deep from the cortical leptomeninges into the brain parenchyma. From there, CSF and solutes may re-enter the lymphatic or venous circulation. This CSF circulation has been described under the term of glymphatic system (7, 8). The rapid elimination of both Patent Blue dye and gemcitabine from the brain to the liver and urine in our two experimental models is consistent with this route. Furthermore, we can understand why Patent Blue penetrates better into the grey matter, which is more vascularized by the cortical meningeal arteries, than into the white matter, which is more vascularized by the central arteries.

There are limitations in our conclusions that are based on 2 non-human models of the blood-brain barrier. Moreover, intraventricular route circumvents the blood-brain barrier in two animal models without brain tumors. It seems unlikely that the barrier is radically different in humans and sheep, but this is difficult to prove clinically in patients. Similarly, we did not investigate the impact of ventricular perfusion on the distribution of Patent Blue V or gemcitabine from the cerebrospinal fluid (CSF) to an existing brain tumor. This investigation could be interesting in a xenograft model in mice or nude rats. Unfortunately, there is no experimental brain tumor model in large animals such as sheep where continuous 24-hour intraventricular perfusion is easier than in rats or mice.

We selected GEM for the treatment of brain tumors for several reasons. GEM is an antimetabolite that does not induce necrotic toxicity in normal central nervous system cells and has previously been administered intrathecally or intracerebrally in both animals and humans (9, 10, 11, 12). GEM has been evaluated preclinically in medulloblastoma, a pediatric brain tumor that can be difficult to treat in certain subtypes. It has been used in combination with pemetrexed (13), axitinib (14), or ribociclib (15) in in vitro models of neurospheres and in vivo xenografts. Its activity is considered promising in these models. Gemcitabine (GEM) has moved from pre-clinical into clinical trials for children with medulloblastoma as part of the St Jude Children’s Research Hospital study SJMB12 (now closed). Intravenous GEM plus pemetrexed was assigned to Group 3 Medulloblastoma patients. Study results are pending (ClinicalTrials.gov:ID NCT01878617). Moreover, GEM displays a broad anticancer spectrum that could be interesting to treat meningeal metastasis from solid tumors if cytotoxic concentration is achieved in brain and CSF. Data independently retrieved from the Cancer Cell Line Encyclopedia (CCLE) show that prolonged exposure (72 h) to nanomolar concentrations of GEM inhibits the growth of most human cancer cell types (16, 17). In CCLE, GEM IC50 was inferior to 100 nM (26 µg/l) for most of breast, lung adenocarcinoma and melanoma cell lines, human tumors that are most often involved in meningeal metastases. In CCLE, GEM is also active against human glioma cell lines, including low-grade gliomas and GB, with IC50 values ranging from 10 to 1000 nM (2.6–260 µg/L). In our *in vitro* experiments, the mean IC50 was 7.3 µg/L and the IC90 was approximately 100 nM (26 µg/L) after a 24-hour exposure. Residual cells beyond the IC90 came into a state of growth arrest suggestive of senescence (18, 19). As not all the biochemical markers of a definitive senescence are present in the residual cells (< 10 % by definition), resumption of cell proliferation should be the subject of further examination

A limitation of our study is the restricted number of glioma models used, limited to three cell lines, which may not adequately recapitulate the heterogeneity and complexity of glioblastoma. Future investigations should incorporate more advanced and physiologically relevant models, such as patient-derived glioblastoma organoids, especially those enriched for glioma stem cell (GSC)-like populations, to better capture tumor architecture, cellular diversity, and invasive properties.

Our results and those from CCLE indicate that glioma cells are not intrinsically resistant to GEM; rather, the BBB limits its clinical efficacy. We further showed that prolonged exposure to GEM over 24 hours is more effective than a 1-hour administration, even at higher concentrations, consistent with its cell cycle–dependent activity during the S phase (20). Finally, GEM demonstrated greater *in vitro* activity against GB cells than cytarabine, a related antimetabolite previously used intrathecally for carcinomatous meningitis (21). Given the lower efficacy of cytarabine and its lack of availability, GEM appears more suitable for intrathecal approaches in GB.

Because prolonged exposure to GEM is more cytotoxic toward cancer cells than short 1-hour infusion and may bypass the blood–brain barrier, we investigated a 24-hour intraventricular GEM infusion protocol in sheep, a large animal model with body weight close to that of humans. Based on our *in vitro* data, we defined a minimal therapeutic target corresponding to the IC90 for glioma cells and lung, breast and melanoma cell lines, i.e. GEM concentrations ≥ 26 µg/L in cerebrospinal fluid (CSF) or ≥ 26 µg/kg in brain tissue. At the end of a 24-hour intraventricular infusion of 20 mg GEM in sheep, GEM concentrations reached 1 415 µg/L in CSF and 850 µg/kg in brain tissue. The elevated concentrations achieved *in vivo*, relative to active *in vitro* concentrations, supports the potential for antitumoral activity in gliomas.

Only a few studies have examined the pharmacology and tolerability of intrathecal GEM. In non-human primates, weekly intrathecal injections of 5 mg GEM for four weeks resulted in peak ventricular CSF concentrations of 297 ± 105 µg/mL, with a short half-life; concentrations fell below 0.03 µg/mL after 6 hours, and transient CSF pleocytosis was the only reported toxicity (9). GEM uptake was studied in glioblastoma in phase 0 study (21). GEM dose was 500 or 1000 mg/m2 just before surgery to 10 GBM patients, who were biopsied after 1-4 h. Tumor gemcitabine level varied from 60 to 3580 pmol/g tissue (15 to 920 µg/kg). In our experiments, GEM concentrations reached mean of 850 µg/kg in normal brain tissue of sheep after a prolonged time (24 h). The area under the curve (AUC) of GEM concentration in brain tissue and CSF would be worth studying in animals between IV injection and prolonged intraventricular perfusion. Unfortunately, there is no model of brain tumor in big animal to investigate the blood to tumor barrier. Our observations indicate that prolonged intraventricular GEM infusion could be more effective than the veinous route. We believe that the broad anti-tumor spectrum of GEM, when administered for a prolonged period of 24 hours, should be investigated in brain tumor models in animals, and then ultimately in humans, children or adults, particularly when a surgical re-intervention is performed for a recurrent glioma or medulloblastoma.

Clinical safety data are limited to a single phase I study in patients with carcinomatous meningitis (12). Ten patients received intrathecal GEM bolus injections. A weekly 5 mg dose was tolerated in three patients; however, dose escalation was associated with reversible neurotoxicity (transverse myelitis at 5 mg bi-weekly and somnolence at 10 mg bi-weekly). The estimated CSF half-life was approximately 50 minutes. No objective responses were observed, and further development of intrathecal GEM was discontinued. Only one case report has described clinical efficacy of intrathecal GEM bolus administration in meningeal metastases from lung adenocarcinoma (23).

Here, we observed that GEM doses of 5, 10, and 20 mg were well tolerated in sheep when administered as a 24-hour intraventricular infusion. Further investigation could be carried out in the sheep model. The maximum tolerated dose (MTD) was not reached at 20 mg in sheep, indicating that even higher intraventricular dose may warrant further investigation. Similarly, the tolerability of repeated intraventricular GEM infusions, i.e. administered weekly, should be evaluated in the sheep model prior to clinical translation. As intraventricular GEM could be considered as an adjuvant therapy following surgical resection of recurrent glioblastoma, medulloblastoma, or brain metastases. The safety of ventricular GEM administration after lobectomy also requires dedicated evaluation.

In conclusion, given the pharmacological advantage of prolonged 24-hour exposure of gemcitabine (GEM) to cancer cells, the ability to bypass the blood–brain barrier via intraventricular infusion, and the favorable tolerability observed in animal models, we propose that new phase I clinical trials of intraventricular GEM should be considered for refractory glioma, medulloblastoma or meningeal metastasis from solid tumors. Furthermore, the sheep intraventricular infusion model may serve as a valuable preclinical platform to evaluate other conventional or emerging anticancer agents.

## Acknowledgments

We thank Pr Florence LEFRANC, Pr François DUCRAY, Dr Anthony FICHTEN, Pr Luc TAILLANDIER, Pr Alain BRON, and Dr Louis LARROUQUERE for their helpful discussion. We thank Claire DABONNEVILLE, Dr François MAISONNEUVE, Dr Reda GARIDI, Dr Séverine FRITOT for their technical help and Dr. Zuzana SAIDAK for reviewing the English version. In memoriam Pr François MARTIN.

